# Endothelial c-Src mediates neovascular tuft formation in Oxygen-Induced Retinopathy

**DOI:** 10.1101/2024.04.17.589627

**Authors:** Emmanuelle Frampton, Priyanka Som, Brittany Hill, Alexander Yu, Ivar Noordstra, Emma Gordon, Lilian Schimmel

**Affiliations:** Centre for Cell Biology of Chronic Disease, Institute for Molecular Bioscience, The University of Queensland, St Lucia, Brisbane, Queensland, Australia

**Keywords:** diabetic retinopathy, neovascularisation, c-Src, vascular tufts

## Abstract

**Introduction:** Vascular retinopathy, characterised by abnormal blood vessel growth in the retina, frequently results in vision impairment or loss. Neovascular tufts, a distinctive pathological feature of this condition, are highly leaky blood vessel structures, exacerbating secondary complications. Despite their clinical significance, the mechanisms underlying tuft development are not fully elucidated, posing challenges for effective management and treatment of vascular retinopathy. In this study, we investigate the role of c-Src in neovascular tuft formation. Although c-Src has been acknowledged as a pivotal regulator in developmental angiogenesis within the retinal vasculature, its specific role in governing pathological retinal angiogenesis remains to be fully understood.

**Methods:** We utilised the Oxygen-Induced Retinopathy (OIR) model to induce the formation of neovascular tufts in both Cre-mediated vascular specific c-Src knockout mice and their wildtype littermates. Subsequently, we conducted high-resolution imaging and analysis of isolated retinas, to elucidate the precise role of c-Src in the formation of vascular tufts.

**Results:** c-Src depletion demonstrated a significant reduction in the formation of neovascular tufts within the OIR model, underscoring the pivotal role of c-Src in pathological retinal angiogenesis. Notably, this decrease in tuft formation was observed independently of any alterations in cell death, cell proliferation or cell adhesion and the absence of c-Src did not impact tuft pericyte coverage and junctional morphology.

**Conclusion:** These findings underscore the critical role of c-Src in the pathogenesis of neovascular tufts in vascular retinopathy. Understanding the molecular mechanisms involving c-Src may offer valuable insights for the development of targeted therapies aimed at mitigating vision-threatening complications associated with retinopathy.

## Introduction

Diabetes mellitus affects around 422 million people worldwide and is strongly associated with cardiovascular diseases. Within the diabetic population, vascular complications pose a prevalent risk, particularly those affecting the delicate microvasculature of the retina, leading to the development of retinopathies. Notably, diabetic retinopathy stands as one of the most prevalent ocular pathologies impacting around 35.4% of diabetic patients worldwide [1]. The progression of diabetic retinopathy unfolds through two distinct phases. In the initial ‘non-proliferative’ stage, elevated blood glucose levels inflict damage upon the vascular capillaries, giving rise to microaneurysms, haemorrhages and ischemic conditions [2]. Subsequently, the onset of ischemia triggers the production of pro-angiogenic factors such as Vascular Endothelial Growth Factor (VEGF), inducing the advanced ‘proliferative’ stage. Here, abnormal blood vessels, known as neovascular tufts, proliferate on the retinal surface, exhibiting a characteristic knot-like morphology [3]. The impaired permeability of these tufts often leads to increased fluid accumulation, exacerbating complications like retinal detachment and ultimately culminating in blindness [4,5].

The discovery that VEGF contributes to pathological neovascularisation and disease progression prompted the development of VEGF and VEGF Receptor 2 (VEGFR2) inhibitors that have been utilised in the clinic to treat patients with proliferative retinopathies [6,7].

Nonetheless, up to 40% of patients display a sub-optimal treatment response or undesirable off-target effects, demonstrating the necessity for a deeper comprehension of key mechanisms [8,9]. In this regard, adopting a more focussed strategy targeting downstream effectors of VEGF signalling holds promise for potentially achieving greater efficacy with significantly reduced side effects.

Among these effectors, c-Src has emerged as a compelling candidate. This non-receptor tyrosine kinase orchestrates a range of cellular processes crucial to angiogenesis, including cell motility, cell survival, cell adhesion and vascular permeability [10–13]. While c-Src has been recognised as a crucial regulator for developmental angiogenesis in the retinal vasculature, the role of c-Src in controlling pathological retinal angiogenesis is still unclear [14].

Previous research has demonstrated that VEGF-induced activation of c-Src leads to the phosphorylation of the cell-cell junctional protein VE-cadherin [12,15]. Specifically, in mice, Phosphorylation of VEGFR2 at tyrosine 949 by VEGF facilitates the binding of the T-cell-specific adapter (TSAd) protein, a crucial step for the activation of c-Src at cell-cell junctions [16]. While TSAd lacks intrinsic kinase activity, it functions as a scaffold to recruit c-Src to cell junctions. Once activated, c-Src phosphorylates VE-cadherin, triggering its internalisation and the formation of microscopically visible intercellular gaps. This cascade of VEGFR2-TSAd-c-Src signalling ultimately leads to VEGF-induced vascular permeability in the context of developmental angiogenesis [17]. Perturbation of the VEGFR2-TSAd-c-Src signalling axis at either the VEGFR2 level, by mutating Y949, or at the TSAd level, via EC-specific knockdown, leads to a marked reduction in neovascular tuft formation in a mouse model of oxygen induced retinopathy (OIR), emphasising the importance of this pathway in driving of pathological tuft formation [18]. Intriguingly, despite a reduction in neovascularisation, tufts of comparable size exhibit reduced permeability in mutated Y949-mutated VEGFR2 retinas compared to wildtype counterparts, revealing the uncoupling of vascular permeability from angiogenesis under pathological conditions. Furthermore, the Y949 VEGFR2 mutation does not elicit alterations in activated c-Src levels, suggesting that while c-Src might not directly regulate tuft permeability, it may still be involved in pathological angiogenesis [18].

Work on the role of c-Src in developmental angiogenesis revealed that c-Src mediates vessel stability via cell-matrix interactions rather than VE-cadherin based cell-cell adhesions [14]. Focal adhesions establish cell-matrix interactions, and activation of c-Src, either upstream or downstream of Focal Adhesion Kinase (FAK), leads to paxillin phosphorylation and enforcement of the adhesion [19]. Disruption of paxillin phosphorylation by c-Src deficient endothelial cells destabilises the cell-matrix connections and causes vascular regression in mice [14].

The potential involvement of c-Src in both the formation and permeability of retinal neovascular tufts remains elusive, presenting a critical gap in our understanding of pathological angiogenesis. In this study, we address this knowledge gap by leveraging the Oxygen-Induced Retinopathy (OIR) model to dissect the role of c-Src in the intricate process of pathological neovascularisation within the retina.

## Methods

### Mice

Animals were cared for and kept at the UQBR facility under laminar airflow at 22-25°C and with a 12-hour light/dark cycle. All animal work conducted was approved by the University of Queensland’s Molecular Biosciences Animal Ethics Committee (permits IMB/231/17/BREED, IMB/161/20/BREED, IMB/174/23/BREED, IMB/424/17, IMB/349/20 and IMB/543/23). Inducible endothelial specific deletion of c-Src (*c-Src*^*fl/fl*^*;Cdh5-CreERT2*) was generated by crossing *c-Src*^*fl/fl*^ mice with tamoxifen inducible Cre recombinase (*Cdh5CreERT2)* [14]. C57BL6 (*wt*) mice were received from the Animal Resources Centre, Perth, Australia and used for outcross with *c-Src*^*fl/fl*^*;Cdh5-CreERT2* in order to generate *Cdh5-CreERT2* mice.

### Genotyping

DNA for genotyping was extracted from mouse samples with Quick Extract (QE09050, Astral Scientific). PCR using GoTaq (M123, Promega) was performed to genotype for presence of *CreERT2* and c-Src deletion. Genotyping for CreERT2 was performed using forward primer: CTGACCGTACACCAAAATTTGCCTG and reverse primer: GATAATCGCGAACATCTTCAGGTT. To verify successful deletion of c-Src, genotyping of P15 mice was performed using forward primer: GGCGTTGTAAGAGCACT and reverse primer: CTAGTCCTCTGAGAGAGGCTGAG. PCR bands were observed in 2% agarose gel and imaged using GelDoc XR+ (Biorad).

### Inducible gene deletion

Cre activity and c-Src gene deletion were induced by intraperitoneal injections of pups (male and female) with 400 μg tamoxifen (Sigma T5648) at P10, P11 and P12. Mice were sacrificed at P15.

### Oxygen-Induced Retinopathy (OIR)

Adapted from the oxygen induced retinopathy protocol described in Connor et al., [20]. Mice were kept in normoxia (21% oxygen) from P0 to P7. On P7, the mice litter along with the lactating adult female, was placed into the oxygen chamber (BioSpherix/Laftech) and oxygen in the chamber is maintained at 75% for three consecutive days: P7, P8 and P9. The lactating female mouse was given a one-hour break on P8 and P9, to allow her to de-stress. A maximum of 6 pups on one female adult were placed inside the chamber for improved survival of the litter. On P10, the litter and the lactating female were taken out of the oxygen chamber and returned to 21% oxygen [21]. Tamoxifen driven gene deletion was induced at P10, P11 and P12, during neovascularisation. Mice were sacrificed at P15.

### Retina immunohistochemistry (IHC)

At P15, eyes were collected and fixed for 20 minutes in 4% paraformaldehyde (PFA) at room temperature. The eyes were washed in phosphate-buffered saline (PBS) and retinas were dissected. The retinas were blocked for 2 hours at room temperature in 1% FBS (Gibco), 3% BSA (Sigma), 0.5% Triton X-100 (Sigma), 0.01% Sodium deoxycholate (Sigma), 0.02% Sodium Azide (Sigma) in PBS at pH 7.4. Samples were incubated overnight at 4°C with primary antibodies in blocking buffer, followed by 5 washes in PBS and incubation with the appropriate secondary antibody for 2 hours at room temperature in blocking buffer. After 5 washes in PBS, retinas were post-fixed in 4% PFA for 3 minutes at room temperature before mounting in Prolong Gold antifade medium (Invitrogen). Images were acquired using a Zeiss LSM710 and Zeiss LSM880 confocal microscope. Z-stacks and tile-scans were used for whole retina imaging with Plan 10x/0.45 Air objective. High magnification images of neovascular tufts were acquired using 40x/1.1 Water or 63x/1.15 Water objectives.

### Antibodies

The following primary antibodies were used (IHC 1:200); rat anti-mouse VE-cadherin (Becton Dickinson, 555289), rabbit anti-mouse Cleaved Caspase-3 (Asp175) (Cell Signalling, 9661), rabbit anti-Ki67 (abcam, 15580), rabbit anti-phospho-paxillin(Y118) (Invitrogen, 44-722G), rabbit anti-NG2 (Millipore, AB5320). Isolectin B4 directly conjugated to Alexa 488, 568 and 647 (ThermoFisher Scientific, I21411, I21412, I32450) (IHC 1:500). NucBlue™ Live ReadyProbes™ Reagent (Hoechst 33342) (ThermoFisher Scientific, R37605) (IHC 2 drops/ml). Fluorescently labelled secondary antibodies were obtained from Invitrogen (IHC 1:250)

### Retina image analysis and quantification

#### Avascular area quantification

FIJI was used to stitch whole retina tile-scans. Percentage avascular area was measured as described by Connor et al. [20] using Adobe Photoshop. In brief, the lasso tool was used to manually outline the retina and measure total retina area. A separate selection was made to measure the avascular areas per retina. The percentage avascular area of total retina area was calculated. All measurements were normalised to the average of the wildtype mice within the same litter.

#### Neovascular tuft area quantification

FIJI was used to stitch whole retina tile-scans. Percentage tuft area was measured as described by Connor et al. [20] using Adobe Photoshop. In brief, the lasso tool was used to manually outline all tufts within the retina. Total tuft area was measured and percentage tuft area of total vascular area (using total retina area minus avascular area) was calculated. All measurements were normalised to the average of the wildtype mice within the same litter.

#### Cell death analysis

Analysis of Cleaved Caspase-3, cell death marker, was performed using FIJI. For each image, the tuft area was manually defined using Isolectin B4 staining to create a ROI. Within the tuft ROI the number of cells expressing Cleaved Caspase-3 were counted and normalised to the total area of the respective tuft ROI. All measurements were normalised to the average of the wildtype mice within the same litter.

#### Cell proliferation analysis

Analysis of Ki-67, cell proliferation marker, was performed using IMARIS. For each image, the tuft area was manually defined using Isolectin B4 staining to create a 3D ROI. Within the tuft ROI the number nuclei (NucBlue) were counted and the number of Ki-67-positive cells were counted. The percentage of Ki-67-positive cells of total number of cells was normalised to the total area of the tuft ROI. All measurements were normalised to the average of the wildtype mice within the same litter.

#### Phosphorylated paxillin analysis

Analysis of phosphorylated (p)-paxillin)-paxillin^Y118^ was performed using IMARIS. For each image, the tuft area was manually defined using Isolectin B4 staining to create a 3D ROI. Within the tuft ROI the Sum Intensity of p-paxillin ^Y188^ was measured and normalized to the total area of the respective ROI. To account for variations in staining intensity between samples, p-paxillin ^Y188^ signal in two non-vascular ROIs was measured, averaged as the Sum Intensity per area, and used to normalise the tuft p-paxillin ^Y188^. Three tuft images per retina were quantified and averaged to determine p-paxillin intensity per retina (n = one retina). All measurements were normalised to the average of the wildtype mice within the same litter.

#### Pericyte coverage analysis

Analysis of NG2 was performed using FIJI. For each image, the tuft area was manually defined using Isolectin B4 staining to create a ROI. Within the tuft ROI the NG2 signal was thresholded, pericyte area was measured and percentage to the total area of the respective tuft ROI was calculated. All measurements were normalised to the average of the wildtype mice within the same litter.

#### VE-cadherin pattern classification

Pattern classification for junctional VE-cadherin was performed using a blinded image analysis approach (Bentley et al., 2014). For each image, the tuft area was manually defined using Isolectin B4 staining to create a ROI. Within the tuft ROI the VE-cadherin signal was max projected and images were randomised to prevent biased comparison between *c-Src*^*fl/fl*^ and *c-Src*^*fl/fl*^*;Cdh5-CreERT2* tufts. These randomised images were further processed into 16×16 μm patches. The VE-cadherin morphology in each patch was manually classified as ‘active’ (serrated, discontinuous and internal vesicular pattern), ‘intermediate’ (continuous but serrated pattern), or ‘inactive’ (straight continuous pattern). Total number of patches per retina was used to determine percentile of each pattern.

### Statistical analysis

Generation of graphs and all statistical analysis was performed using GraphPad Prism. All data are represented as mean ± SEM. Specification on statistical tests and sample size is reported in the figure legends. Only significant comparisons are displayed in the graphs.

## Results

### Endothelial specific c-Src deletion reduces OIR-induced neovascular tuft formation

Oxygen-Induced Retinopathy (OIR) is an experimental model used to study retinal neovascularisation [20,21]. A schematic of the experimental set up is shown in Fig 1A. After normal vascular development during the first 6 postnatal days (Fig 1B), mice are exposed to a high level of oxygen (hyperoxia) from postnatal day 7 to 10. This hyperoxia initiates disruption of the normal development of the retinal vasculature, leading to vaso-obliteration or regression of existing blood vessels compared to mice not subjected to OIR (Fig 1C-D). Following the hyperoxic phase, the mice are returned to room air, which induces a relative hypoxic state in the retina due to the sudden decrease in oxygen levels. In response to hypoxia, high levels of VEGF are produced and initiate a process of pathological angiogenesis characterised by the formation of abnormal, leaky blood vessels, referred to as neovascular tufts. Neovascular tufts are manifested as a knot-like structure with rounded edges comprised of endothelial cells which lack phenotypical sprouting protrusions (Fig 1E). This dynamic process ultimately results in a disorganised, dysfunctional vasculature characteristic of pathological conditions such as in diabetic retinopathy.

**Figure 1.**
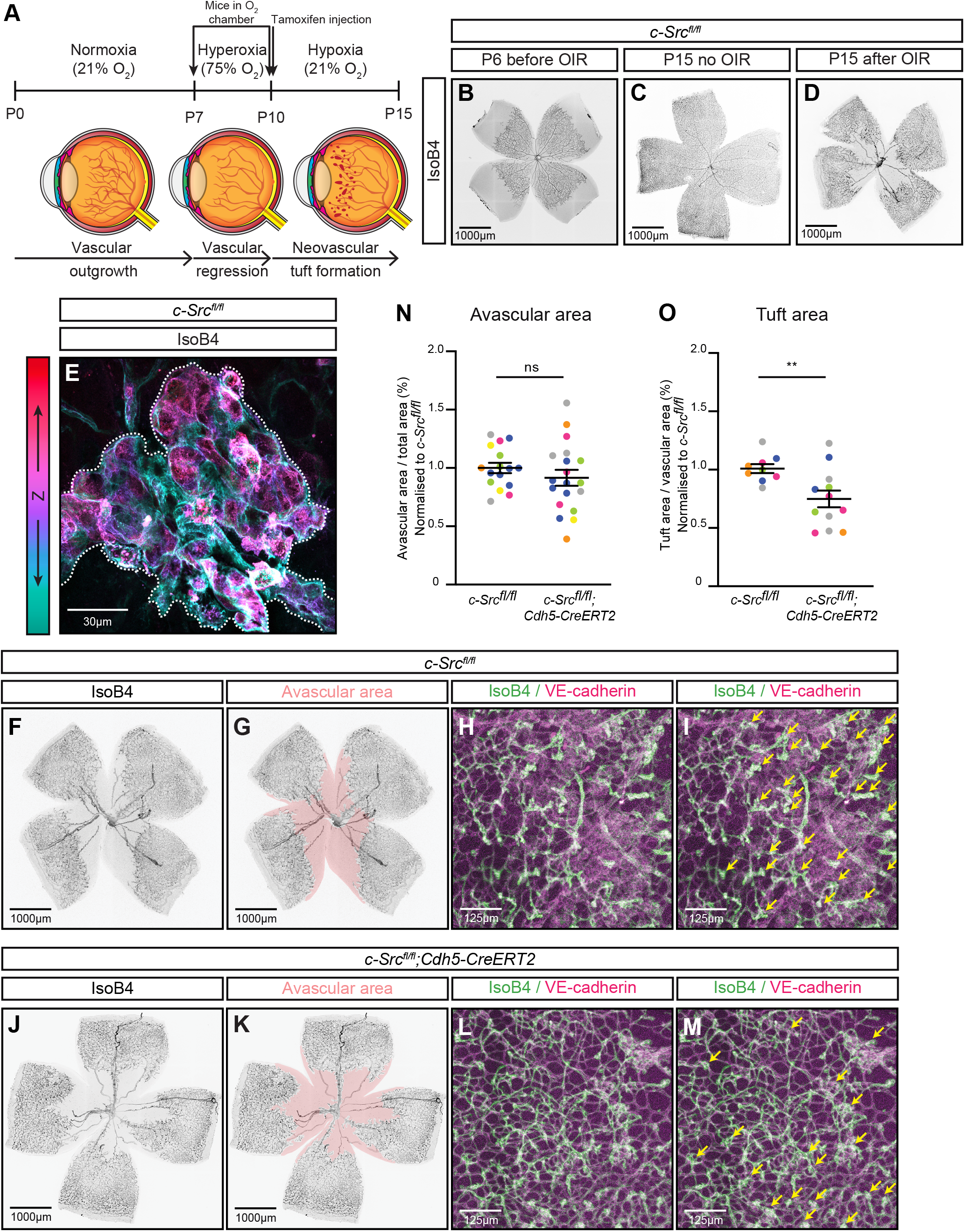
Endothelial specific c-Src deletion reduces OIR-induced neovascular tuft formation. **(A)** Schematic of the OIR model showing timeline of experiment with mice placed in the oxygen chamber at P7 to induce vascular regression. Mice are removed from the oxygen chamber and injected with tamoxifen to initiate c-Src deletion. Normal oxygen levels result in induction of hypoxia and neovascular tuft formation. Mice are sacrificed at P15. **(B)** Representative image of *c-Src*^*fl/fl*^ retina at P6 in a normal developing mouse. Immunofluorescent staining was performed for Isolectin B4 (IsoB4, grey). **(C)** Representative image of *c-Src*^*fl/fl*^ retina at P15 in normal developing mouse. Immunofluorescent staining was performed for Isolectin B4 (IsoB4, grey). **(D)** Representative image of c-Srcfl/fl retina at P15 in OIR subjected mouse. Immunofluorescent staining was performed for Isolectin B4 (IsoB4, grey). **(E)** High magnification image of representative neovascular tuft of OIR subjected *c-Src*^*fl/fl*^ mouse. Immunofluorescent staining was performed for Isolectin B4 (IsoB4, colour code indicates Z position). Tuft area is outlined with white dashed line. **(F)** Representative image of OIR subjected *c-Src*^*fl/fl*^ retina. Immunofluorescent staining was performed for Isolectin B4 (IsoB4, grey). **(G)** Representative image of avascular area (pink) in OIR subjected *c-Src*^*fl/fl*^ retina. **(H)** Representative image of neovascular tufts in OIR subjected *c-Src*^*fl/fl*^ retina with immunofluorescent staining for Isolecting B4 (IsoB4, green) and VE-cadherin (magenta). **(I)** Indication of neovascular tufts (yellow arrows) in OIR subjected *c-Src*^*fl/fl*^ retina with immunofluorescent staining for Isolecting B4 (IsoB4, green) and VE-cadherin (magenta). **(J)** Representative image of OIR subjected *c-Src*^*fl/fl*^*;Cdh5-CreERT2* retina. Immunofluorescent staining was performed for Isolectin B4 (IsoB4, grey). **(K)** Representative image of avascular area (pink) in OIR subjected *c-Src*^*fl/fl*^*;Cdh5-CreERT2* retina. **(L)** Representative image of neovascular tufts in OIR subjected *c-Src*^*fl/fl*^*;Cdh5-CreERT2* retina with immunofluorescent staining for Isolecting B4 (IsoB4, green) and VE-cadherin (magenta). **(M)** Indication of neovascular tufts (yellow arrows) in OIR subjected *c-Src*^*fl/fl*^*;Cdh5-CreERT2* retina with immunofluorescent staining for Isolecting B4 (IsoB4, green) and VE-cadherin (magenta). **(N)** Quantification of percentage avascular area normalised to the average of the wildtype mice within the same litter. n= 14-19 retinas from 6 litters. **(O)** Quantification of percentage tuft area normalised to the average of the wildtype mice within the same litter. n= 9-12 retinas from 5 litters All data are represented as mean ±SEM with individual data points indicated and colours represent different litters. Statistical significance was determined using Mann-Whitney test (N, O) ^**^ p<0.01.

Under pathological conditions of OIR, research has demonstrated that blocking the phosphorylation of VEGFR2 on Y949 effectively reduces the formation of neovascular tufts [18]. The activation of the VEGFR2-TSAd-c-Src pathway has been identified as crucial for angiogenic sprouting [12]. Notably, the absence of endothelial c-Src results in diminished retinal angiogenesis during development [14]. Therefore, the requirement of c-Src in the formation of OIR-induced neovascular tuft formation was investigated. To specifically address the requirement for c-Src in the formation of neovascular tufts, endothelial c-Src deletion was induced after the hyperoxic phase via tamoxifen injections from P10 (Fig 1A). Tamoxifen injection of *c-Src*^*fl/fl*^*;Cdh5-CreERT2* mice results in c-Src protein deletion (Schimmel et al., 2020) and Cre activity upon tamoxifen injection for each experiment was verified by PCR of P15 mice (Sup Fig 1A). As expected, the OIR-induced avascular area in endothelial specific c-Src deleted mice (*c-Src*^*fl/fl*^*;Cdh5-CreERT2*) remained unchanged when compared to Cre-negative wildtype littermates (*c-Src*^*fl/fl*^) (Figure 1F-G, J-K, N). However, the total neovascular tuft area was significantly decreased in c-Src deleted retinas compared to wildtypes, indicated by Isolectin B4 and VE-Cadherin staining (Figure 1H-I, L-M, O).

Tamoxifen-activated CreERT2 has previously been demonstrated to influence retinal angiogenesis during development, irrespective of gene deletion [22]. To ensure the observed reduction in tuft formation in c-Src deleted mice was not attributed to interference from tamoxifen-activated CreERT2, OIR subjected retinas from wildtype C57Bl6 mice (*wt*) were compared to CreERT2 outcross mice (*Cdh5-CreERT2*), revealing that activation of CreERT2 by tamoxifen has no effect on OIR induced vascular tuft formation (Sup Fig 1B-K). This underscores the essential role of c-Src activation in OIR-induced neovascular tuft formation.

### Neovascular tuft reduction is independent of apoptosis, proliferation, or vessel-matrix adhesion

As we observed a reduction in neovascular tuft area in *c-Src*^*fl/fl*^*;Cdh5-CreERT2* retinas (Fig 1O), we reasoned this could be due to alterations in cell death, cell proliferation or vascular instability. Endothelial apoptosis rates in neovascular tufts was not altered in c-Src depleted (*c-Src*^*fl/fl*^*;Cdh5-CreERT2*) mice compared to wildtype littermates (*c-Src*^*fl/fl*^) indicated by Cleaved Caspase-3 staining (Fig 2 A-C). Furthermore, we did not observe significant changes in the percentage of Ki-67-positive, proliferating cells within neovascular tufts (Fig 2 D-F).

**Figure 2.**
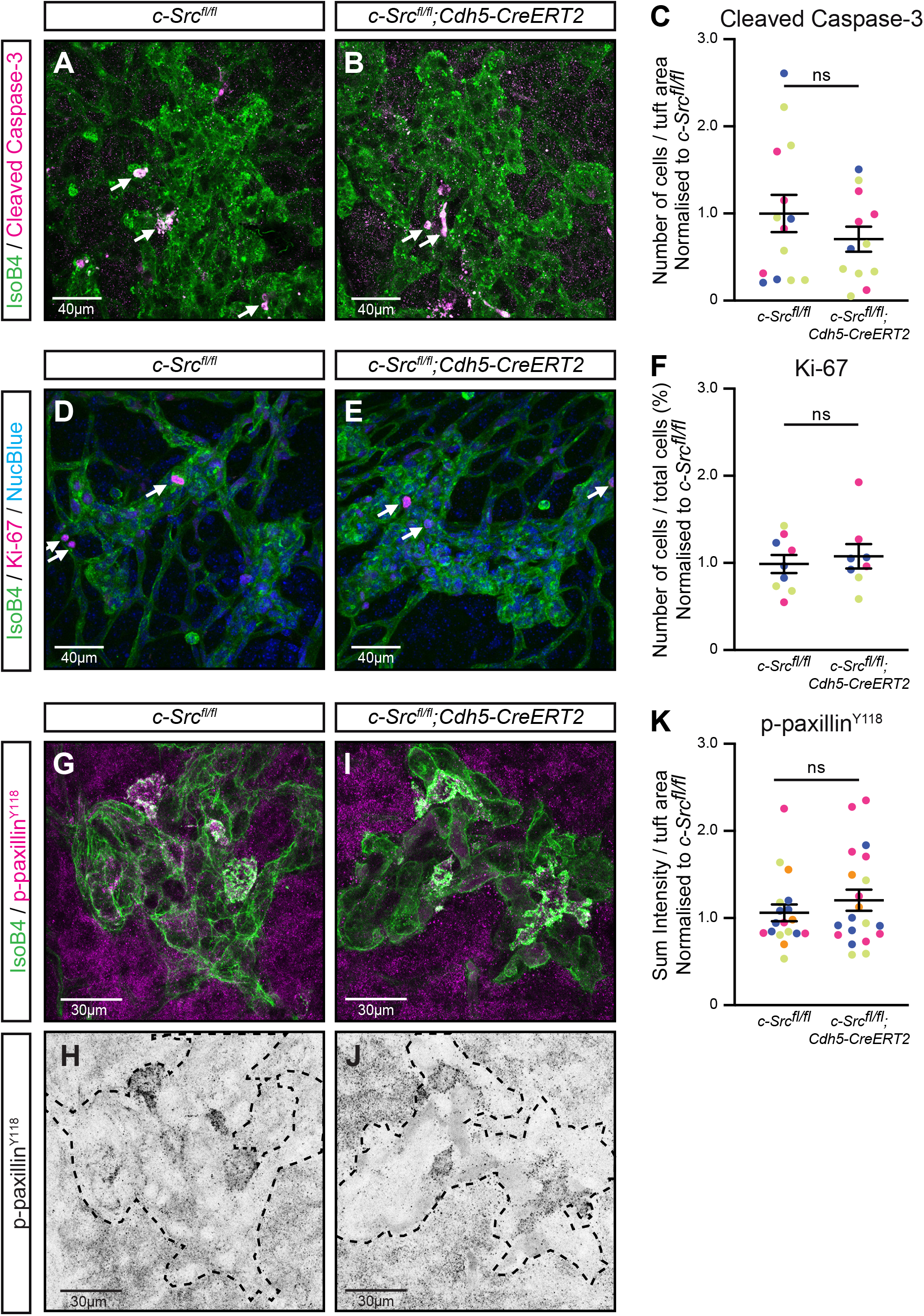
Neovascular tuft reduction is independent of apoptosis, proliferation, or vessel-matrix adhesion. **(A)** Representative image of OIR subjected *c-Src*^*fl/fl*^ tuft with immunofluorescent staining for Isolectin B4 (IsoB4, green) and Cleaved Caspase-3 (Magenta). **(B)** Representative image of OIR subjected *c-Src*^*fl/fl*^*;Cdh5-CreERT2* tuft with immunofluorescent staining for Isolectin B4 (IsoB4, green) and Cleaved Caspase-3 (Magenta). White arrows indicate Cleaved Caspase 3 positive cells. (**C)** Quantification of number of Cleaved Caspase-3 positive cells per neovascular tuft area, normalised to the average of the wildtype mice within the same litter. n=12-14 retinas from 3 litters, with each data point representing the average of 4-5 measured tufts within the same retina. **(D)** Representative image of OIR subjected *c-Src*^*fl/fl*^ tuft with immunofluorescent staining for Isolectin B4 (IsoB4, green), proliferation marker (Ki-67, magenta) and nuclei (NucBlue, blue). **(E)** Representative image of OIR subjected *c-Src*^*fl/fl*^*;Cdh5-CreERT2* tuft with immunofluorescent staining for Isolectin B4 (IsoB4, green), proliferation marker (Ki-67, magenta) and nuclei (NucBlue, blue). White arrows indicate Ki-67 positive nuclei. (**F)** Quantification of percentage Ki-67 positive cells of total cell number within a neovascular tuft, corrected for tuft area and normalised to the average of the wildtype mice within the same litter. n=8-9 retinas from 3 litters, with each data point representing the average of 4-5 measured tufts within the same retina. **(G)** Representative image of OIR subjected *c-Src*^*fl/fl*^ tuft with immunofluorescent staining for Isolectin B4 (IsoB4, green) and p-paxillin^Y118^ (Magenta). **(H)** Black and white image of p-paxillin^Y118^ signal only of image shown in (G). **(I)** Representative image of OIR subjected *c-Src*^*fl/fl*^*;Cdh5-CreERT2* tuft with immunofluorescent staining for Isolectin B4 (IsoB4, green) and p-paxillin^Y118^ (Magenta). **(J)** Black and white image of p-paxillinY118 signal only of image shown in (I). Tuft area is outlined with black dashed line. (**K)** Quantification of p-paxillin^Y118^ Sum Intensity per neovascular tuft area corrected for background signal intensity and normalised to the average of the wildtype mice within the same litter. n=18-20 retinas from 4 litters, with each data point representing the average of 4-5 measured tufts within the same retina. All data are represented as mean ±SEM with individual data points indicated and colours represent different litters. Statistical significance was determined using Mann-Whitney test (C, F, K).

These findings align with previous work indicating that c-Src does not regulate cell proliferation nor cell death during developmental retinal angiogenesis [14]. Overall, our results suggest that c-Src is not involved in mediating cell apoptosis or proliferation during neovascular tuft formation, and that reduction of tufts in c-Src depleted mice likely involves a different mechanism.

In addition to cell apoptosis or proliferation, vessel stability is strongly dependent on endothelial cell attachment to the surrounding matrix. Vessel-matrix interaction, facilitated via focal adhesions, are essential for vessel stability and prevent vessel regression. We previously showed that during development, c-Src-deficient endothelial cells of the retina displayed significantly reduced phosphorylated (p)-paxillin^Y118^, causing an increase in vascular regression [14]. To further investigate the regulation of focal adhesions in mediating c-Src-dependent stabilisation in pathological neovascular tufts, we examined p-paxillin^Y118^ in OIR retinas. Immunofluorescent staining for p-paxillin^Y118^ revealed no significant difference in tufts of c*-Src*^*fl/fl*^*;Cdh5-CreERT2* mice compared to *c-Src*^*fl/fl*^ mice (Fig 2G-K), indicating that endothelial specific c-Src deletion does not alter vessel-matrix adhesion in neovascular tufts.

### Stabilisation and junction morphology of neovascular tufts is not mediated by c-Src

Pericytes tightly wrap around vessels providing stability and limiting permeability [23,24]. This interaction is affected in diabetic retinopathies by secretion of platelet-derived growth factor-B (PDGF-B) by endothelial cells, resulting in abnormal pericytes at neovascular tufts increased vascular permeability [25,26]. Depletion of pericytes in OIR models results in impaired revascularisation and reduces neovascular tuft formation [26]. Examining the coverage of neovascular tufts by pericytes could provide insight into the observed reduction in tuft formation in c-Src depleted retinas. However, immunofluorescent staining of OIR subjected retinas for pericyte marker NG2, revealed no significant change in NG2-positive pericyte coverage of tufts in *c-Src*^*fl/fl*^*;Cdh5-CreERT2* retinas compared to wildtype littermates (*c-Src*^*fl/fl*^) (Fig 3A-E). This suggests that c-Src may not be directly involved in regulating vessel stability through pericyte recruitment during pathological angiogenesis.

**Figure 3.**
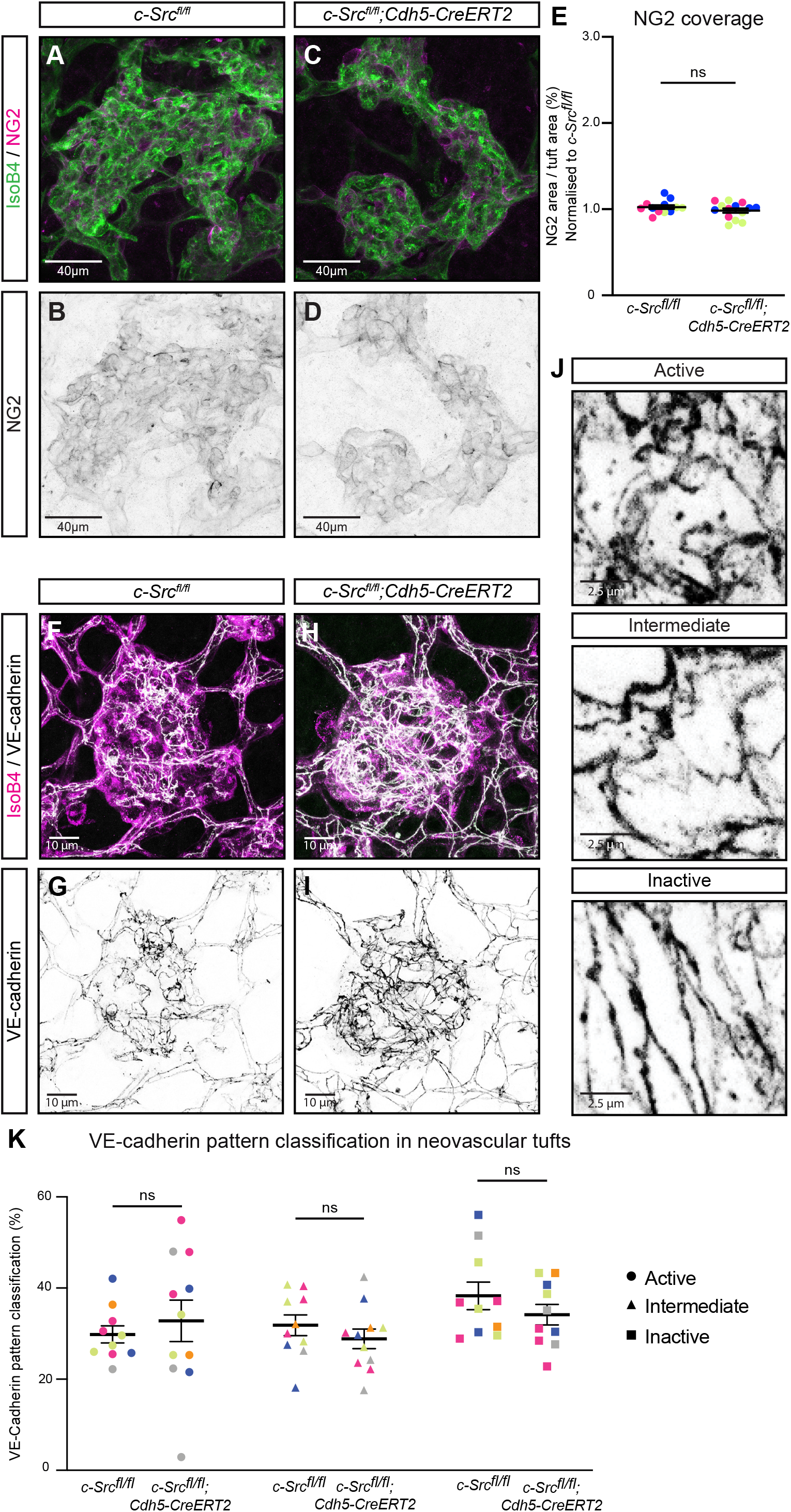
Stabilisation and junction morphology of neovascular tufts is not mediated by c-Src. **(A)** Representative image of OIR subjected *c-Src*^*fl/fl*^ tuft with immunofluorescent staining for Isolectin B4 (IsoB4, green) and pericytes (NG2, Magenta). **(B)** Black and white image of NG2 signal only of image shown in (A). **(C)** Representative image of OIR subjected *c-Src*^*fl/fl*^*;Cdh5-CreERT2* tuft with immunofluorescent staining for Isolectin B4 (IsoB4, green) and pericytes (NG2, Magenta). **(D)** Black and white image of NG2 signal only of image shown in (C). (**E)** Quantification of percentage NG2 area of total neovascular tuft area per and normalised to the average of the wildtype mice within the same litter. n=12-14 retinas from 3 litters, with each data point representing the average of 4-5 measured tufts within the same retina. **(F)** Representative image of OIR subjected *c-Src*^*fl/fl*^ tuft with immunofluorescent staining for Isolectin B4 (IsoB4, magenta) and VE-cadherin (white). **(G)** Black and white image of VE-cadherin signal only of image shown in (F). **(H)** Representative image of OIR subjected *c-Src*^*fl/fl*^*;Cdh5-CreERT2* tuft with immunofluorescent staining for Isolectin B4 (IsoB4, magenta) and VE-cadherin (white). **(D)** Black and white image of VE-cadherin signal only of image shown in (H). **(J)** Examples of VE-cadherin pattern classifications as ‘active’ (serrated, discontinuous, and internal vesicular pattern), ‘intermediate’ (continuous but tortuous pattern), and ‘inactive’ (straight, continuous pattern). (**K)** Quantification of percentage VE-cadherin pattern classification in neovascular tufts. n=10-11 mice from 5 litters, with each data point representing the average of 1-2 measured retinas per mouse. All data are represented as mean ±SEM with individual data points indicated and colours represent different litters. Statistical significance was determined using Mann-Whitney test (E) or Kruskal Wallis test (K).

During angiogenesis, endothelial cells underdo dynamic rearrangements to mediate sprout elongation. These rearrangements are mediated by differential cell-cell adhesion based on VE-cadherin localisation [27]. The internalisation of VE-cadherin following phosphorylation facilitates junctional dynamics, allowing for endothelial cell rearrangement within developing sprouts. However, this mechanism is compromised in retinopathy [27]. To assess whether loss of c-Src leads to alterations in endothelial junction morphology, we utilised quantitative high-resolution confocal image analysis on VE-cadherin junctional patterning [27]. VE-cadherin patterning was analysed specifically in neovascular tuft areas of *c-Src*^*fl/fl*^*;Cdh5-CreERT2* mice compared to wildtype littermates (*c-Src*^*fl/fl*^) (Fig 3F-I). Segmented VE-cadherin junctions were classified based on a gradient between ‘active’ (serrated, discontinuous, and internal vesicular pattern), ‘intermediate’ (continuous but tortuous pattern), and ‘inactive’ (straight, continuous pattern) (Fig 3J). No changes in VE-cadherin patterning were observed between wildtype and c-Src-deficient tufts (Fig 3K), suggesting that c-Src does not regulate differential VE-cadherin junctional dynamics in neovascular tufts.

Taken together, endothelial specific c-Src depletion reduces the formation of neovascular tufts in an oxygen induced retinopathy model. The decrease in tuft formation was observed independently of alterations in cell death, cell proliferation or cell adhesion and the absence of c-Src did not impact pericyte coverage or the morphology of the cell-cell junctions within tufts.

## Discussion

Our work sheds light on the critical role of c-Src in the intricate process of pathological angiogenesis, exemplified in conditions such as diabetic retinopathy, but also applicable to wet age-related macular degeneration and retinopathy of prematurity [28]. Notably, c-Src emerges as a key driver in the formation of neovascular tufts. Of particular significance is the discovery that targeted deletion of endothelial c-Src during the proliferative stage yields a substantial reduction in neovascular tuft formation. This finding underscores the therapeutic potential of modulating c-Src activity to mitigate the deleterious effects of aberrant retinal angiogenesis. Despite the marked reduction in neovascular tuft formation following endothelial c-Src deletion, no discernible changes were observed in cell death, cell proliferation or cell adhesion. While this leaves the precise mechanism for c-Src-mediated neovascular tuft formation elusive, it also underscores the complexity of the underlying processes, inviting further investigation and deeper exploration into the intricacies of angiogenic regulation.

Neovascular tufts are notorious for their significant leakage [18], often linked to a loss of pericytes [25,29]. While we demonstrated that c-Src deletion does not change the rate of pericyte coverage in neovascular tufts, delving deeper into whether c-Src mediates permeability could yield valuable insights. Vascular permeability is regulated by VE-cadherin based cell-cell junctions, and phosphorylation-induced internalisation of VE-cadherin results in increased vessel leakage [17]. In OIR retinas, VE-cadherin becomes phosphorylated on multiple residues, leading to the disruption of cell-cell junctions [18]. However, the involvement of c-Src in directly phosphorylating VE-cadherin in neovascularisation is questionable. Perturbation of the VEGF-VEGFR2-TSAd pathway by mutating Y949 in VEGFR2 reduces vascular permeability in OIR retinas but did not show any changes in c-Src activation, indicating that c-Src is not activated downstream of VEGFR2 in the specific setting of OIR [18]. Additional research directly investigating the mechanisms responsible for VE-cadherin phosphorylation and vascular permeability showed that the Src-family kinase Yes, rather than c-Src, is responsible for VE-cadherin phosphorylation [30]. Those findings are supported by our results, revealing that the specific deletion of endothelial c-Src had no discernible impact on the morphology of VE-cadherin. Although VE-cadherin morphology correlates with cell-cell junction integrity, it only serves as a proxy for vascular permeability. Future experiments to directly assess integrity, such as extravasation of fluorescent dyes, leukocytes, or red blood cells, will offer additional insights into the regulation of retinal vascular permeability by c-Src.

c-Src plays pivotal roles in a plethora of cellular processes, encompassing proliferation, differentiation, survival, adhesion, and migration [31,32]. These functions collectively offer potential mechanisms by which the deletion of c-Src could lead to diminished formation of neovascular tufts. We demonstrated that c-Src depletion does not alter proliferation or apoptosis. Given that neovascular tufts observed in retinopathies lack the typical phenotypic formation of vascular sprouts [3], it implies a deficiency in guidance and migration towards the growth factor cue VEGF. The observed decrease in neovascular tuft formation in c-Src-depleted retinas may stem from alterations in both cell adhesion and migration, which are intricately linked to the essential cytoskeletal component, actin. Indeed, actin emerges as a compelling candidate, having been demonstrated to be regulated by c-Src through a number of mechanisms [31].

Firstly, c-Src can directly phosphorylate actin-binding proteins to regulate actin dynamics. For instance, its binding and phosphorylation of cortactin enhances actin filament assembly [33]. Conversely, c-Src mediates the phosphorylation of cofilin, an actin-binding molecule that promotes filament severing and depolymerisation. This leads to its rapid ubiquitination and subsequent proteasomal degradation of cofilin, resulting in actin stabilisation [34].

Secondly, c-Src has been shown to interact with and regulate the activity of Rho family GTPases, which are well established mediators of the cytoskeleton [31,35]. It can directly phosphorylate Rac1, a process implicated in the regulation of the actin cytoskeleton dynamics during cell migration [36]. Additionally, c-Src can phosphorylate proteins involved in the regulation of Rho GTPases. For example, c-Src-mediated phosphorylation of RhoGDI, which typically sequesters inactive GDP-bound Rho GTPases in the cytosol, disrupts the Rho GTPase-RhoGDI complex. This leads to the release and activation of Rho GTPases such as RhoA, Rac1 or Cdc42 mediated actin polymerisation, branching or contraction [36,37]. The variety of interactions with actin-binding proteins and Rho family GTPases allows c-Src to fine tune actin dynamics.

Finally, c-Src plays a crucial role in regulating focal adhesions, which are essential for mediating cell adhesion through interactions with the extracellular matrix [19]. Our findings indicate that c-Src deletion does not cause changes in the focal adhesion component p-paxillin^Y118^ compared to wild type littermate tufts. However, focal adhesions are dynamic structures. Therefore, future investigations should explore the implications of c-Src deletion on focal adhesion dynamics and the potential impact on neovascular tuft formation.

Moreover, the interplay between focal adhesions and cell-cell junctions facilitates coupling of these two compartments via the actin cytoskeleton [36]. Focal adhesion turnover can result in the direct phosphorylation of β-catenin [38], leading to the dissociation of the VE-cadherin-catenin complex and subsequent induction of vascular permeability [39]. This underscores that c-Src-mediated regulation of focal adhesions not only controls cell-matrix adhesion, but through crosstalk via the actin cytoskeleton, it indirectly influences vascular permeability.

In conclusion, c-Src emerges as a central regulator in the intricate process of OIR-induced vascular tuft formation. Its multifaceted roles in modulating actin dynamics, cell adhesion and signalling pathways underscore its significance in orchestrating this complex phenomenon. However, while our current understanding provides valuable insights, further investigations are essential to unravel the precise molecular mechanisms through which c-Src governs neovascular tuft formation. By delving deeper into these mechanisms, we can pave the way for novel therapeutic strategies aiming at mitigating pathological angiogenesis and its associated complications in disease.

## Statements

## Acknowledgements

We thank Lena Claesson-Welsh, Deb Barkauskas and Nicholas Condon for helpful discussions. Microscopy was performed at the Institute for Molecular Bioscience Microscopy Facility which was established with the support of the Australian Cancer Research Foundation (ACRF) and incorporates the Dynamic Imaging, Cancer Biology Imaging and Cancer Ultrastructure and Function Facilities.

## Author contributions

Conceptualisation: E.G., L.S.; Methodology and validation: E.F., P.S., B.H., A.Y., E.G.; Formal analysis: E.F., P.S.; Data curation: E.F., P.S., B.H.; A.Y. Writing - original draft: I.N., L.S.; Visualisation: E.F., I.N., L.S.; Supervision: E.G., L.S.; Project administration: E.F., E.G.; Funding acquisition: E.G., L.S.

## Data availability statement

All data are available in the main text or the supplementary materials.

