## Supplementary Figure 1 for "Endothelial c-Src mediates neovascular tuft formation in Oxygen-Induced Retinopathy"

### Supplemental Figure 1

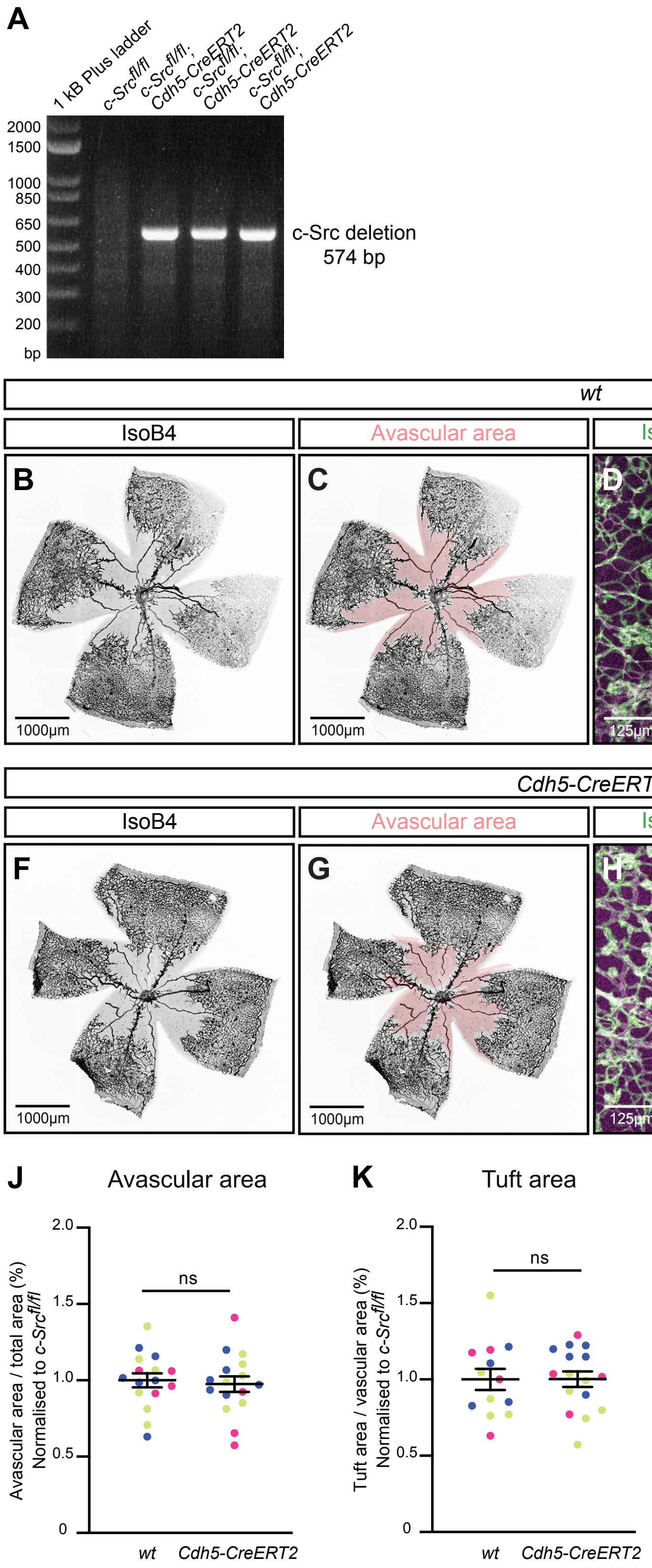

**Supplementary Figure 1. Tamoxifen-activated CreERT activity has no effect on OIR-induced neovascular tuft formation.**

**(A)** Representative image of DNA gel electrophorese of c-Src deletion PCR on wildtype littermate (*c-Src*$^{fl/fl}$) and three *c-Src*$^{fl/fl}$*;Cdh5-CreERT2* mice on samples taken at P15. **(B)** Representative image of OIR subjected *wt* (C57BL6) retina. Immunofluorescent staining was performed for Isolectin B4 (IsoB4, grey). **(C)** Representative image of avascular area (pink) in OIR subjected *wt* (C57BL6) retina. **(D)** Representative image of neovascular tufts in OIR subjected *wt* (C57BL6) retina with immunofluorescent staining for Isolectin B4 (IsoB4, green) and VE-cadherin (magenta). **(E)** Indication of neovascular tufts (yellow arrows) in OIR subjected *wt* (C57BL6) retina with immunofluorescent staining for Isolectin B4 (IsoB4, green) and VE-cadherin (magenta). **(F)** Representative image of OIR subjected *Cdh5-CreERT2* retina. Immunofluorescent staining was performed for Isolectin B4 (IsoB4, grey). **(G)** Representative image of avascular area (pink) in OIR subjected *Cdh5-CreERT2* retina. **(H)** Representative image of neovascular tufts in OIR subjected *Cdh5-CreERT2* retina with immunofluorescent staining for Isolectin B4 (IsoB4, green) and VE-cadherin (magenta). **(I)** Indication of neovascular tufts (yellow arrows) in OIR subjected *Cdh5-CreERT2* retina with immunofluorescent staining for Isolectin B4 (IsoB4, green) and VE-cadherin (magenta). **(J)** Quantification of percentage avascular area normalised to the average of the wildtype mice within the same litter. n= 16 retinas from 3 litters. **(K)** Quantification of percentage tuft area normalised to the average of the wildtype mice within the same litter. n= 13-16 retinas from 3 litters. All data are represented as mean ±SEM with individual data points indicated and colours represent different litters. Statistical significance was determined using Mann-Whitney test (J, K).
